# Fibromyalgia patients with high levels of anti-satellite glia cell IgG antibodies present with more severe symptoms

**DOI:** 10.1101/2022.07.06.498940

**Authors:** Emerson Krock, Carlos E. Morado-Urbina, Joana Menezes, Matthew A. Hunt, Angelica Sandström, Diana Kadetoff, Jeanette Tour, Vivek Verma, Kim Kultima, Lisbet Haglund, Carolina B. Meloto, Luda Diatchenko, Eva Kosek, Camilla I. Svensson

**Author notes:** Athinoula A. Martinos Center for Biomedical Imaging, Massachusetts General Hospital, Harvard Medical School and Department of Radiology, Massachusetts General Hospital, Boston, MA, USA. Shared last authorship (Eva Kosek, Camilla I. Svensson). Corresponding author; Address: Karolinska Universitetssjukhuset, Centrum för Molekylär Medicin, L8:03, 17176 Stockholm.

## Abstract

**Objective:** Transferring fibromyalgia patient IgG to mice induces pain-like behaviour and fibromyalgia IgG binds mouse and human satellite glia cells (SGCs). These findings suggest that autoantibodies could be part of fibromyalgia pathology. However, it is unknown how frequently fibromyalgia patients have anti-SGC antibodies and how anti-SGC antibodies associate with disease severity.

**Methods:** We quantified serum or plasma anti-SGC IgG levels in two fibromyalgia cohorts from Sweden and Canada using an indirect immunofluorescence murine cell culture assay. Fibromyalgia serum IgG binding to human SGCs in human dorsal root ganglia tissue sections was assessed by immunofluorescence (n=14/group).

**Results:** In the cell culture assay anti-SGC IgG levels were increased in both fibromyalgia cohorts compared to controls. Elevated anti-SGC IgG was associated with higher levels of self-reported pain in both cohorts, and higher fibromyalgia impact questionnaire scores and increased pressure sensitivity in the Swedish cohort. Anti-SGC IgG levels were not associated with fibromyalgia duration. Swedish FM patients were clustered into FM-severe and FM-mild groups and the FM-severe group had elevated anti-SGC IgG compared to the FM-mild and controls. Anti-SGC IgG levels detected in culture were positively correlated with increased binding to human SGCs. Moreover, the FM-severe group had elevated IgG binding to human SGCs compared to the FM-mild and control groups.

**Conclusions:** A subset of fibromyalgia patients have elevated levels of anti-SGC antibodies, and the antibodies are associated with more severe fibromyalgia severity. Screening fibromyalgia patients for anti-SGC antibodies could provide a path to personalized treatment options that target autoantibodies and autoantibody production.

## INTRODUCTION

Fibromyalgia (FM) is a prototypical condition of nociplastic pain and despite a prevalence of 2-4%, the pathogenesis is unclear. FM is characterized by chronic widespread pain, hypersensitivity to touch and temperature, fatigue, cognitive difficulties, elevated nociceptive neuron sensitivity and activity, and reduced intraepidermal nerve fiber density (1–6). Due to the unclear pathogenesis of FM, treatment options have unsatisfactory efficacy and diagnostic tests do not exist. In the case of FM, as with other nociplastic pain conditions, i.e., conditions characterized by altered nociception that is not fully explained by nociceptive or neuropathic pain mechanisms (7,8), a key question is what causes the altered nociception.

To identify potential drivers of nociplastic pain in FM we used a passive transfer mouse model to investigate a role for pathogenic, pain-driving IgG antibodies (9). IgG can induce pain independent of overt inflammation or nerve damage (10–12) and a role for antibodies has been identified in other nociplastic pain conditions (13–15). We found that transferring FM IgG into mice induces pain-like behaviour, nociceptor sensitization and hyperactivity, and decreased intraepidermal nerve fiber density (9), all of which are characteristics of FM (4,5). Finally, we found that FM IgG accumulated in the dorsal root ganglia (DRG). FM IgG bound satellite glia cells (SGCs) *in vivo, in vitro* and in human DRG tissue sections (9). SGCs envelop sensory neuron soma and are linked to nociceptor activity and pain-like behavior in models of rheumatoid arthritis (16), neuropathic (17–19) and inflammatory pain (20,21). Taken together, these findings suggest that SGC-binding autoantibodies underlie some pathological characteristics of FM.

We previously used single patient IgG preparations from eight individuals or pooled IgG preparations from several individuals. Thus, the frequency of FM patients with autoantibodies, and their association with disease severity, remains unclear. To further establish the clinical relevance of our findings, the aim of the current study was to determine the frequency of SGC autoantibodies in FM patients and the relation between FM autoantibodies and FM symptom severity. As we have found that pooled FM IgG binds cultured SGCs and human DRG sections (9). we used these approaches to evaluate the presence of anti-SGC IgG levels in individual patient and control samples. We found individuals with FM have elevated levels of anti-SGC IgG in two regionally distinct cohorts. Moreover, elevated levels of anti-SGC IgG are associated with worse disease severity.

## MATERIALS AND METHODS

### Study participants

#### Karolinska Institutet fibromyalgia and control subjects

Age and sex-matched serum samples were collected from FM patients or healthy controls (HC) between 2015 and 2017 (Ethical permit number: 2014/1604-31/1) and written consent was received from all individuals. FM patients fulfilled the 1990 and 2011 ACR fibromyalgia criteria. Exclusion criteria were other rheumatic or autoimmune diseases, primary causes of pain other than FM, comorbid severe somatic disorders (neurological, cardiovascular, etc), severe psychiatric disorders requiring treatment, previous heart or brain surgery, substance abuse, medication with anticonvulsants or antidepressants, self-reported claustrophobia, inability to refrain from hypnotics, nonsteroidal anti-inflammatory drugs, or analgesics prior to study participation, specifically 48 hours before the visit, hypertension (>160/90 mmHg), obesity (body mass index > 35), smoking (>5 cigarettes/day), magnetic implants, pregnancy, and the inability to understand and speak Swedish. HC subjects were screened by telephone interview, met the exclusion criteria for FM and were free from any chronic pain conditions.

#### McGill University fibromyalgia and control subjects

FM and HC subjects’ plasma was collected at the McGill University rheumatology clinic, the Alan Edwards Pain Management Unit and the Research Institute of the McGill University Health Centre and written consent was received from all participants (Institutional Review Board Study Number A05-M50-14B). Inclusion criteria for all FM subjects were the ability to write and speak English or French, aged 40 or more and a clinical diagnosis of FM by a rheumatologist based on the ACR 2010 criteria. Exclusion criteria were uncontrolled medical or psychiatric conditions, clinical study participation that may interfere with the current study, prior or current drug and/or alcohol abuse and control individuals were additionally excluded if they have been diagnosed with a chronic pain condition or have a history of depression.

#### Osteoarthritis and control subjects

Knee osteoarthritis patients were recruited at the Ortho Center, Upplands Väsby, Sweden and controls were recruited via local newspaper advertisements. Osteoarthritis inclusion criteria were radiographically diagnosed knee osteoarthritis and knee pain as the dominant pain symptom. Patients were excluded if they had chronic pain due to other causes or previous knee surgery. Control subjects had the same exclusion criteria, and their average weekly pain rating was >20 mm VAS. Serum was collected and written consent was received from all patients (local ethical committee approval 2011/2036-31-1).

### Study participant phenotyping

#### Questionnaires

Minimum, maximum, and average global pain intensities over the past week and current pain intensities were assessed using a 100mm VAS. The Fibromyalgia Impact Questionnaire (FIQ), assessing FM-related symptoms and disability, was completed by FM subjects (22). Depressive characteristics were assessed using the Beck Depression Inventory (23). Pain intensity in the McGill cohort was assessed with the global assessment of pain.

#### Pressure pain thresholds (PPTs)

PPTs were assessed using a hand-held algometer (Somedic Sales AB) as previously described (24). A probe (1 cm^2^) was applied with steadily increasing pressure of approximately 50 kPa/second. Subjects pressed a button to indicate pain sensation and the value was recorded. PPTs were assessed bilaterally at 4 anatomical sites (supraspinatus muscle, lateral epicondyle, gluteus muscle, and knee medial fat pad) and the average of all 8 assessments was used for subsequent analyses.

#### Conditioned pain modulation (CPM) score

CPM was evaluated with PPTs as the test stimulus and ischemic pain as the conditioning stimulus (Tourniquet test) as previously described (25). A baseline PPT assessment was performed at the quadriceps femoris muscle (right thigh) with the pressure algometer described above. A blood pressure cuff at the participants’ upper left arm was inflated to 200 mmHg and participants lifted a 1 kg weight by extending their wrist until their perceived pain intensity exceed 50 mm on a 0-100 mm VAS scale. With the cuff inflated, the experimenter assessed the PPTs at the participants’ right thigh repetitively, with at least 10 seconds between each assessment, for 4 minutes or until the participants decided to end the procedure (end PPT value). A CPM score was calculated for each participant: end PPT value – PPT baseline)/PPT baseline. Positive scores represent inhibition, negative scores represent pain facilitation.

### Animals

Female BALB/cAnNRj mice (Janvier, France) aged 12-26 weeks were used to establish cell cultures. Mice were housed in IVC GM-500 cages (Techniplast) in a dedicated temperature control animal room with a 12-hour light/dark cycle with ad libitum access to food and water. Ethical approval was received from the Stockholm North Animal Ethics Board (4945-2018).

### Cell culture and immunocytochemistry

DRG-derived, SGC-enriched and neuron-enriched cultures were established as previously described (9,11,26). Briefly, cells were seeded onto Nunc Lab-Tek glass chamber slides. To enrich the SGCs, the nonadherent cells, including neurons, were removed. The non-adherent cells were seeded onto Nunc Lab-Tek II CC2-treated chamber slides to establish SGC-depleted, neuron-enriched cultures. Cells were then allowed to recover for approximately 16 hours. Serum or plasma samples were diluted 1:100 in 37°C complete culture media, passed through a 0.22 μM syringe filter and incubated with the cells for three hours. Cells were then fixed with 4% paraformaldehyde and antibodies against glutamine synthase or protein gene product 9.5 were used to identify SGCs and neurons, respectively. An anti-human IgG antibody detect human IgG. All antibody details are in table 1. Z-stack images were collected using a Zeiss LSM800 confocal microscope operated by LSM ZEN2012 software (Zeiss).

**Table 1.** Antibody details. GS (glutamate synthase), GFAP (Glial fibrillary acidic protein), PGP (protein gene product), AF (Alexa Fluor).

| Primary Antibodies |  |  |  |  |  |
| --- | --- | --- | --- | --- | --- |
| Target | Host | Manufacturer | Catalogue # | Dilution |  |
| GS | Rabbit | Abcam | ab73593 | 1:500 |  |
| GFAP | Rabbit | Dako | Z0334 | 1:1000 |  |
| NF200 | Chicken | Neuromics | CH22104 | 1:1000 |  |
| PGP 9.5 | Rabbit | Cedarlane | CL7756AP | 1:2000 |  |
| Secondary Antibodies |  |  |  |  |  |
| Target | Fluorophore | Host | Manufacturer | Catalogue # | Dilution |
| Rabbit IgG | AF488 | Goat | ThermoFisher | A11008 | 1:300 |
| Rabbit IgG | Cy2 | Donkey | Jackson ImmunoResearch | 711-225-152 | 1:200 |
| Human IgG | AF594 | Goat | ThermoFisher | A11014 | 1:300 |
| Human IgG | Cy3 | Donkey | Jackson ImmunoResearch | 709-165-149 | 1:600 |
| Chicken IgG | Cy5 | Donkey | Jackson ImmunoResearch | 711-175-152 | 1:500 |
| Human IgG (H+L) Fab Fragment | Unconjugated | Goat | Jackson ImmunoResearch | 109-007-003 | 100 µg/ml |

### Human DRG immunofluorescence

FM IgG reactivity against human DRG tissue was tested as previously described (9). A human DRG from a 53-year-old female organ donor without a history of chronic pain was collected following consent from the next of kin (McGill University Health Centre REB 2019-4896). Tissue was fixed on the slides with 4% PFA, blocked with PBS containing 3% normal donkey serum and 0.3% Triton X-100 and then incubated with 100 μg/ml unconjugated anti-human IgG Fab fragments (H+L, Jackson Immunoresearch). Slides were then incubated with FM or HC serum diluted 1:500 in PBS with 1% normal donkey serum and 0.1% Triton X-100. Slides were incubated with anti-human IgG antibodies, incubated with antibodies against GFAP and NF200, then appropriate secondaries, counterstained with DAPI, and cover-slipped with Prolong Gold mounting media.

### Image analysis

Individual cells were identified using Cellpose, a deep-learning neural network (27). Cellpose was run with Python v3.7.9 and the region of interest of each cell were imported into FIJI. Human IgG binding to SGCs and neurons was then assessed in FIJI and the percentage of cells bound and the average integrated density of IgG binding was determined. Human DRG images were analyzed using the drgquant pipeline as described previously (28). Experimenters were blinded to the serum type that cells or tissue were incubated with.

### ELISA

Total IgG titres were quantified using enzyme linked immunosorbent assays (ELISA, BMS2091, Thermofisher Scientific). ELISAs were performed according to manufacturers’ instructions and read with a Spectramax iD3 plate reader (Molecular Devices) and analyzed with Softmax Pro software (Molecular Devices). Plate effects were corrected using a linear mixed-effects model in RStudio v1.4.1106.

### Statistics

All data is presented as mean ± 95% confidence intervals (CI). Differences in the percentage of cells bound by IgG were analyzed by Mann-Whitney test or a Kruskal-Wallis test and Dunn’s post hoc test. IgG intensity binding, IgG titres and phenotypic data was analyzed by two-tailed t-tests or one-way ANOVA and Tukey’s post hoc test. Spearman rank correlations were used to investigate relationships between IgG binding and phenotypic data. Karolinska FM serum samples were clustered into FM-mild and FM-severe groups using a k-means cluster analysis in RStudio v1.4.1106. A p-value <0.05 was considered significant. All data was analyzed in GraphPad Prism v9 except for correlations which were analyzed in RStudio v1.4.1106.

## RESULTS

### FM serum has increased levels of anti-SGC antibodies

To evaluate levels of satellite glia cell (anti-SGC antibodies) or neuron binding IgG in FM serum we established SGC-enriched and neuron enriched cell cultures from mouse DRGs. Live cell cultures were incubated with serum from single FM patients or healthy control (HC) subjects so that IgG could only bind cell-surface antigens. Anti-SGC IgG levels were quantified by the percentage of SGCs bound by human IgG and the binding intensity assessed by the average integrated density. IgG in FM serum bound a higher percentage of SGCs (54.44%, CI [44.7-64.12] vs 31.54%, CI[22.87, 40.2], p=0.0016) and generated a greater immunofluorescence signal intensity compared to HC serum IgG (2.99, CI [1.94, 4.06] vs 1.31, CI [0.41, 2.21], p=0.017, Fig. 1A-C). These results indicate that, on average, FM serum contained higher levels of anti-SGC antibodies. Importantly, total IgG levels did not differ between groups (Fig. 1D). When examining individual data points (Fig. 1B, E) we observed a large variation between individual serum samples suggesting that only a subset of FM patients have autoreactive IgG antibodies.

**Figure 1.**
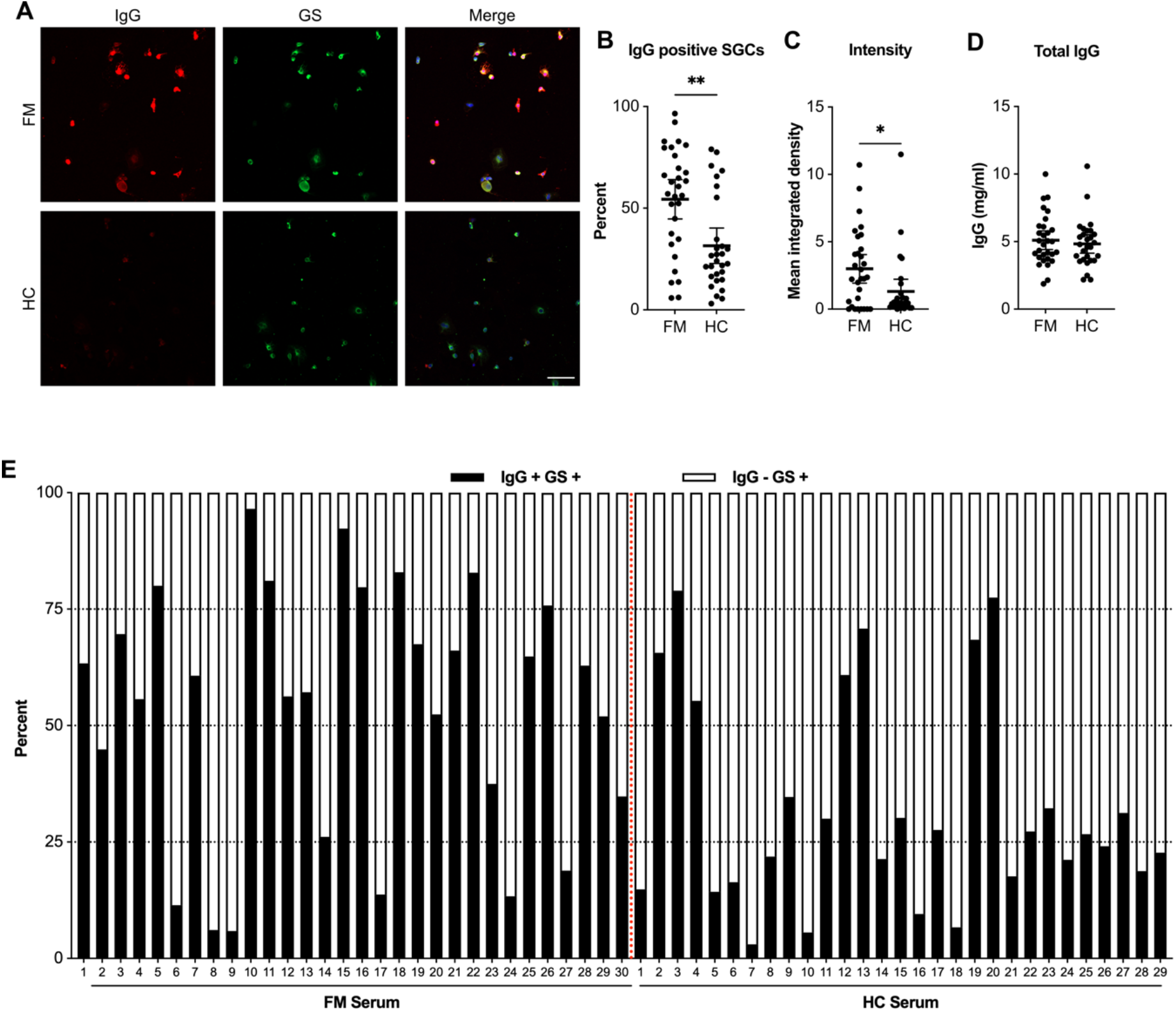
Fibromyalgia serum contains elevated levels of IgG that are reactive with satellite glia cells. Satellite glia cell-enriched cell cultures were incubated with serum from fibromyalgia (FM) or healthy control (HC) individuals and then stained for glutamine synthase (GS), a marker of SGCs. (**A**) Representative images of cultures incubated with FM of healthy control serum where human IgG is red and GS is green. (**B**) The percentage of SGCs that were bound by human IgG was determined for each serum sample tested. (**C**) The integrated density of IgG, as a measure of pixel intensity, was determined for each cell that was imaged and then the mean integrated density was calculated for each serum sample. (**D**) Total IgG serum titres did not differ between groups. (**E**) The percentage of SGCs bound by IgG per donor. Scale bar is 50 μm. N = 30 (FM) or 29 (HC), the difference between the percentage of IgG positive cells was determined with a Mann-Whitney test and the difference between intensity was determined with a two-tailed t-test. * Indicates p<0.05, ** indicates p<0.01.

### FM plasma has increased levels of anti-SGC antibodies in a validation cohort

To validate our findings, we repeated these experiments with a regionally distinct cohort of FM and HC plasma samples from McGill University. We found that FM plasma samples had elevated levels of IgG that bound SGCs, both in terms of IgG+ SGCs (47.65% CI[40.68, 54.61] vs 29.1% CI[22.98, 35.19], p=0.0002) and the binding intensity (1.24, CI[0.90, 1.58] vs 0.83, CI[0.43, 1.23], p=0.0092, Fig. 2A-C). The total IgG levels did not differ in plasma between groups (Fig. 2D). These results indicate that increased levels of anti-SGC antibodies are not specific to a single FM cohort and are present in regionally distinct cohorts.

**Figure 2.**
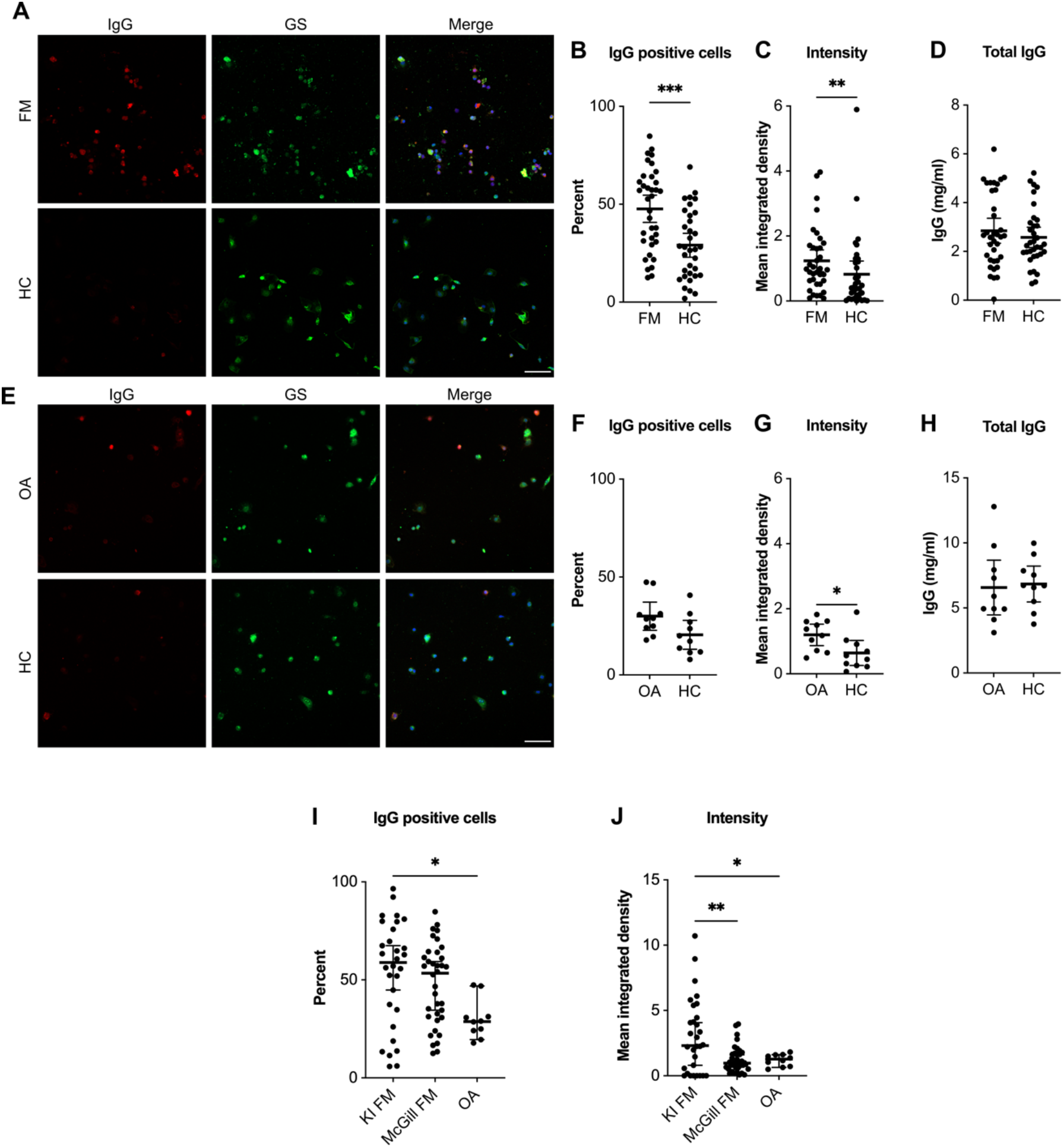
Fibromyalgia plasma contains elevated levels of SGC-reactive IgG in a validation cohort, but osteoarthritis serum does not. SGC-enriched cultures were incubated with plasma collected from a regionally distinct FM cohort. (**A**) Representative images of cultures where human IgG is red and glutamine synthase (GS) is green. (**B**) The percentage of human IgG+ SGCs and (**C**) the integrated density (intensity) of IgG binding were determined. (**D**) Total IgG serum titres did not differ between groups. (**E**) SGC-enriched cultures were incubated with serum from osteoarthritis (OA) and HC individuals. (**F**) The percentage of IgG+ SGCs and (**G**) intensity were determined. (**H**) Total IgG serum titres did not differ between groups. (**I**) The percentage of IgG+ SGCs and (**J**) the binding intensity were compared between the FM and OA cohorts. Scale bar is 50 μm. N= 36 (McGill FM), 34 (McGill HC), 10 (OA and HC) or 30 (Karolinska FM). Percentages were assed using a Mann-Whitney test (**B, F**) or a Kruskal-Wallis test followed by Dunn’s post hoc test (**H**). Intensities were analyzed by two-tailed t-tests (**C, G**) or a one-way ANOVA followed by Tukey’s post hoc test. * indicates p<0.05, ** indicates p<0.01, *** indicates p<0.001.

### Anti-SGC antibodies are not ubiquitous across chronic musculoskeletal pain

To determine if anti-SGC IgG is widespread across chronic musculoskeletal conditions we tested osteoarthritis patient and matched control serum. There was no difference between the percentage of cells bound by IgG in osteoarthritis serum compared to control serum and only a slight elevation of binding intensity (1.2, CI[0.87, 1.53] vs 0.64 CI[0.26, 1.02], p=0.0228, Fig. 2E-G). The total IgG level also did not differ between OA and HC serum.

The Karolinska FM samples bound an elevated level of SGCs compared to the osteoarthritis samples (Fig. 2I) and the binding intensity of the Karolinska samples was greater than the McGill FM samples and the OA samples (Fig. 2J). This further supports that the presence of SGC-reactive IgG is not widespread across chronic pain conditions. However, this comparison also raises the interesting finding that Karolinska FM samples have, on average, a stronger binding profile than the McGill FM samples.

### Anti-neuron IgG levels are not different between FM and controls

Next, we assessed anti-neuron IgG levels. There was no difference in the percentage of neurons bound by IgG nor the binding intensity in the Karolinska FM (Fig. 3A-C), McGill FM (Fig. 3D-F), and osteoarthritis (Fig. 3G-I) samples compared to the matched controls. This suggests that the presence of neuron binding IgG antibodies in FM serum and plasma is not widespread.

**Figure 3.**
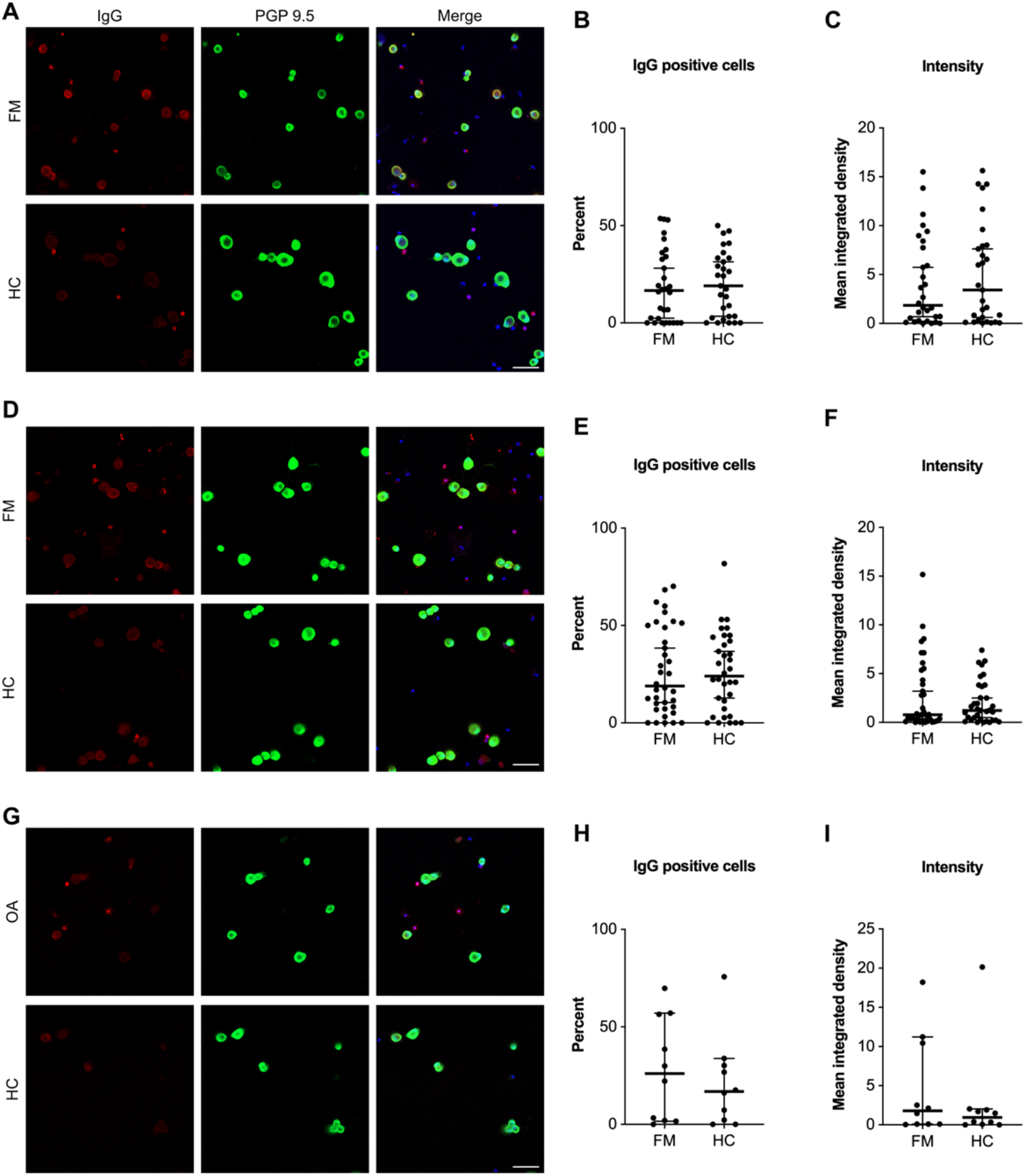
Fibromyalgia and osteoarthritis patients do not have elevated levels of sensory neuron binding IgG. Dorsal root ganglia neuron-enriched/satellite glia cell depleted cultures were incubated with serum from FM patients (**A**), plasma from FM patients (**D**), serum from osteoarthritis (OA) patients or corresponding healthy control individuals (HC). (**A, D, G**) Neurons were identified by PGP 9.5 immunoreactivity (green), IgG binding was detected using an anti-human IgG antibody (red) and cultures were counterstained with Hoechst (blue). The percentage of neurons bound by IgG (**B, E, H**) and the intensity of the binding assessed by integrated density (**C, F, I**) was determined. Scale bar is 50 μm. In **A-C** n=30 (FM) and 29(HC); in **D-F** n=36 (McGill FM), 34 (McGill HC); in **G-H** n=10 (OA and matched HC). Differences in percentages were determined using a Mann-Whitney test and differences in intensity were determined using two-tailed t-tests.

### Anti-SGC antibody levels are correlated with pain intensity

We next investigated whether anti-SGC IgG levels in FM patients correlate with phenotypic characteristics across cohorts. FM patients from McGill were on average older, had a higher BMI and had had FM longer. The maximum pain intensity did not differ between cohorts, but the average pain intensity was greater in Karolinska FM patients compared to McGill FM patients (Fig. 4A-E). Anti-SGC IgG levels, determined by the percentage of cells bound and the binding intensity, were not correlated with age, BMI, total IgG, nor the duration of which an individual has had FM nor chronic pain. However, anti-SGC IgG levels were positively correlated with the self-reported average pain intensity (moderate; IgG+: r=0.62, p<0.0001; Intensity: r=0.61, p<0.0001) and maximum pain intensity (moderate; IgG+: r=0.50, p<0.0001; intensity: r=0.52, p<0.0001) (Fig. 4F). These findings suggest that anti-SGC antibody levels are associated with pain intensity.

**Figure 4.**
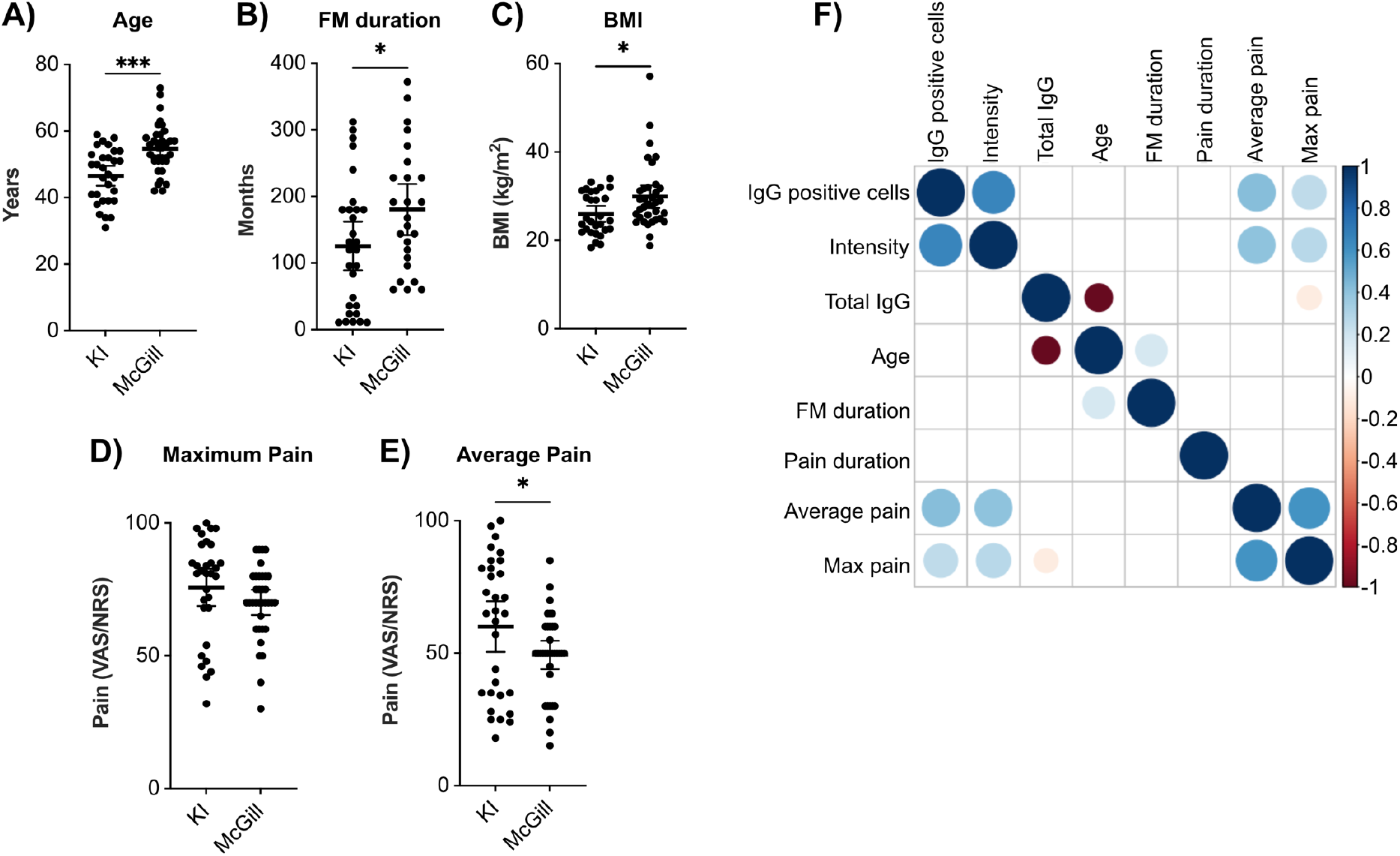
Fibromyalgia IgG binding to satellite glia cells is positively correlated with pain intensity. Comparisons of (**A**) age, (**B**) body mass index (BMI), (**C**) FM duration measured as time from diagnosis, (**D**) maximum pain and (**E**) average pain. Pain intensity was assessed with the visual analogue score of pain at Karolinska and with the numeric rating scale of pain at McGill. Both scales have a range of 0-100. (**F**) Correlations between the percentage of IgG positive cells and the intensity with total IgG levels, age, FM duration, pain duration, and average and maximum pain intensities were examined in FM serum (Karolinska) and plasma (McGill) samples only. The size and colour of the dots indicate the spearman rank correlation coefficient and only statistically significant correlations (p<0.05) have coloured dots whereas non-significant correlations are blank. Differences in age, BMI and pain intensity were assessed by two-way ANOVA followed by Tukey’s post hoc test. Differences in FM duration were analyzed by a two-tailed t-test. * indicates p<0.05, *** indicates p<0.001. N= 30 (Karolinska FM), 36 (McGill FM), 29 (Karolinska HC), 34 (McGill HC) or 66 (FM correlations).

### FIQ, pain thresholds and intensity are correlated with anti-SGC IgG levels

Karolinska FM patients had additional phenotypic characterization, allowing for further analysis. When analyzing the Karolinska FM cohort alone anti-SGC IgG levels were positively correlated with the average, current, minimum and maximum pain intensities (moderate). Additionally, the percentage of cells bound by IgG was positively correlated with the FIQ score (moderate, IgG+: r=0.47, p=0.009) and negatively correlated with an individual’s pressure pain threshold (weak, IgG+: r=-0.37, p=0.046). Of note, anti-SGC IgG levels were not correlated with CPM (Fig. 5A). These correlations indicate that worse disease severity is associated with elevated levels of anti-SGC antibodies.

**Figure 5.**
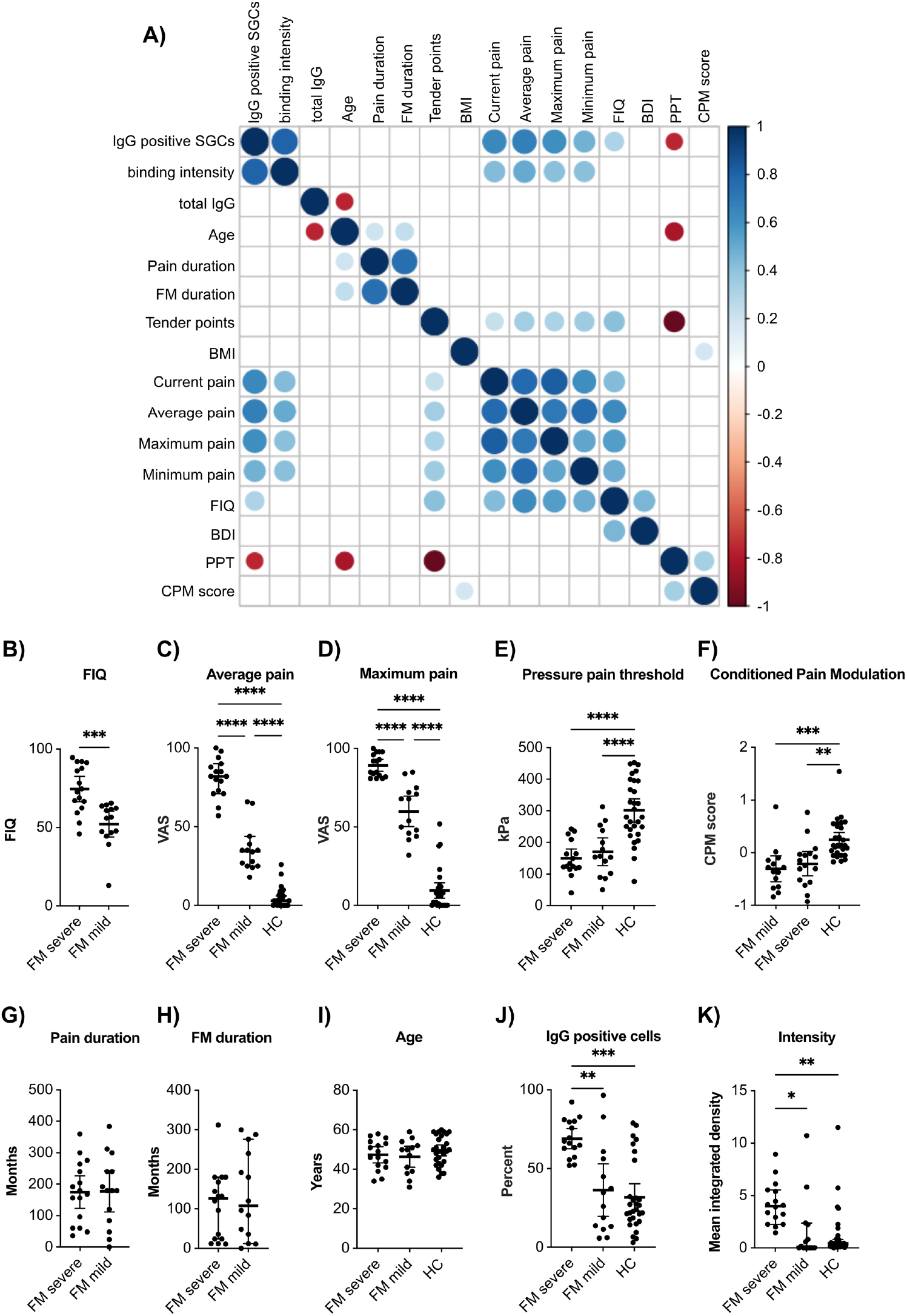
Level of satellite glia cell-reactive fibromyalgia IgG is associated with disease severity. (**A**) The Karolinska cohort has expanded phenotypic data, therefore the correlations between IgG positive cells and binding intensity with various characteristics were examined in the FM samples. The size and colour of the dots indicate the Spearman rank correlation coefficient and only statistically significant correlations (p<0.05) have coloured dots whereas non-significant correlations are blank. The Karolinska FM cohort was divided into two groups based on FIQ, average pain intensity and maximum pain intensity using a k-means cluster analysis. The cluster analysis resulted in a FM severe and a FM mild group (**B-H**). (**J**) The percentage of satellite glia cells bound by IgG and (**K**) the binding intensity were both greater in the FM severe group compared to the FM mild group and the healthy control group (HC). Body mass index (BMI); Fibromyalgia Impact Questionnaire (FIQ); Becks Depression Inventory (BDI); pressure pain threshold (PPT); conditioned pain modulation (CPM). One-way ANOVAs with Tukey’s post hoc test (**C-F**, **I-K**) or two-tailed t-tests (**B, G, H**) were used to examine differences between groups. * indicates p<0.05, ** indicates p<0.01, *** indicates p<0.001.

### Individuals with more severe FM have elevated levels of anti-SGC IgG

The Karolinska FM cohort was split into FM-mild and FM-severe groups using a k-means cluster analysis with the average and maximum pain intensities and the FIQ score. The FM-severe group had elevated FIQ scores and pain intensities compared to the mild group and HC (Fig. 5B-D). Pressure pain thresholds, condition pain modulation scores, chronic pain duration, FM duration and age did not differ between FM groups (Fig. 5E-I). The FM-severe group (IgG+: 68.96%, CI[62.58, 75.32]; Intensity: 4.21, CI[3.12, 5.29]) had an increased percentage of IgG-bound SGCs and the binding intensity was greater compared to the FM-mild group (IgG+: 36.36%, CI[19.74, 52.99), p=0.003; intensity: 1.621, CI[-0.16, 3.39], p=0.0166) and the HC group (IgG+: 31.8%, CI[23.23, 40.38], p=0.0001; intensity: 1.31, CI[0.41, 2.21], p=0.0012; Fig. 5K, J). However, the FM-mild group did not differ from HC. These results suggest that SGC-reactive IgG levels are elevated in FM patients with more severe disease.

### FM serum has elevated levels of IgG that binds human SGCs

Finally, we wanted to determine whether FM serum samples with high levels of anti-SGC IgG also bound human DRG tissue. Human DRG tissue sections were incubated with FM serum containing high or low levels of anti-SGC IgG or HC serum detected in culture. SGC-bound IgG intensity (3.65, CI[3.47, 3.82]) was greater in the serum samples with high levels of anti-SGC IgG compared to serum with low levels of anti-SGC IgG (3.27, CI[3.002, 3.54], p=0.033) and controls (3.22, CI[3.02, 3.41], p=0.012), whereas there was no difference between serum with low anti-SGC IgG and HC serum (Fig. 6A, B). There was also no difference in IgG binding to neuronal soma, axons and non-SGC, non-neuronal tissue (Fig. 6C-E). The IgG binding intensity to human SGCs was moderately correlated with anti-SGC IgG levels determined in SGC-enriched cultures (IgG+: r=0.43, p=0.004; Intensity: r=0.41, p=0.006), but there was no association with total IgG levels or age (Fig. 6F). Finally, IgG binding intensity to human SGCs in the FM-severe group (3.66, CI[3.5, 3.83]) was elevated compared to the FM-mild group (3.26, CI[2.99, 3.52], p=0.014) and HC group (3.18, CI[3.0, 3.26], p=0.003Fig. 6G). Together, these findings indicate that serum from patients with severe FM has elevated levels of antibodies that bind human SGCs and mouse SGCs.

**Figure 6.**
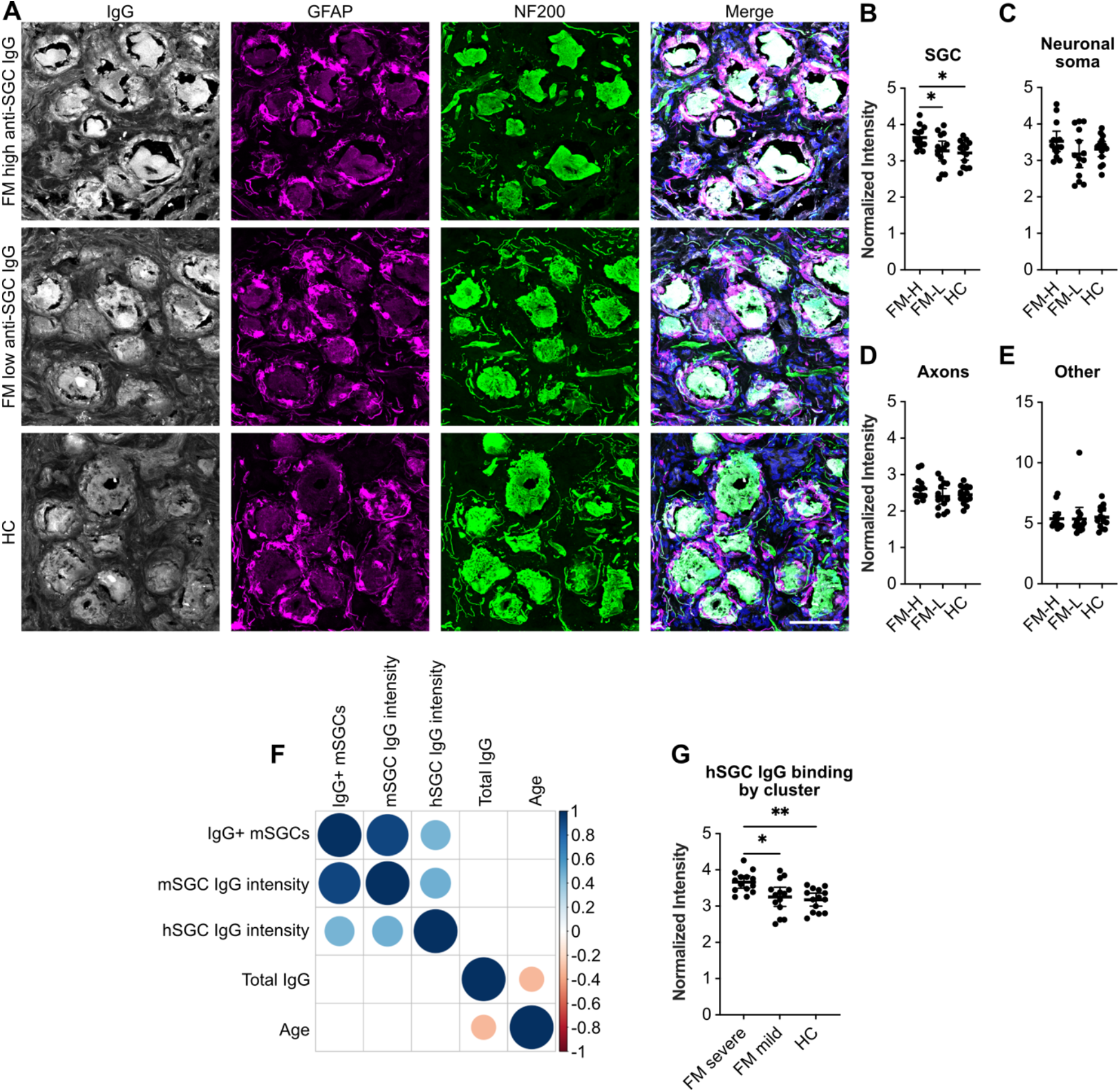
Elevated IgG binding to human satellite glia cells and neurons. (**A**) Human DRG tissue sections were incubated with serum samples that had high levels or low levels of anti-SGC antibodies detected in mouse DRG derived SGC cultures. Human SGCs (hSGCs) were identified with an antibody against glial fibrillary acidic protein (GFAP) and neurons were identified with an antibody against NF200. The normalized pixel intensity of serum IgG binding to (**B**) SGCs, (**C**) neuronal soma, (**D**) axons, and (**E**) GFAP- and NF200-objects was analyzed. (**F**) The intensity of IgG binding to hSGCs correlated with the level of anti-SGC antibodies determined in cell culture by the percent of IgG+ mouse SGCs (mSGCs) and the intensity of that binding. The size and colour of the dots indicate the Spearman rank correlation coefficient and only statistically significant correlations (p<0.05) have coloured dots whereas non-significant correlations are blank. (**G**) Individuals in the FM severe group had elevated IgG binding to hSGCs compared to the FM mild group and HC group. One-way ANOVAs with Tukey’s post hoc test was used to determine differences, * indicates p<0.05, ** indicates p<0.01, n=14/group. Scale bar is 50 μm.

## DISCUSSION

We previously found IgG from FM patients induces FM-like characteristics in mice and that FM IgG binds mouse and human SGCs. However, the frequency of FM patients with SGC binding IgG and the correlation with clinical symptoms had not been examined. Here we found elevated anti-SGC antibodies in two distinct FM cohorts. Moreover, the binding of IgG to SGCs in our cell culture assay correlated with disease severity, indicating the clinical relevance of our finding. Importantly, IgG binding to human SGCs is also elevated compared to controls and correlated with anti-SGC IgG levels detected in culture. FM IgG binding to human SGCs was also elevated in more severe FM, further supporting a link between anti-SGC autoantibodies and disease severity. Furthermore, these results suggest that pathogenic autoantibodies either underlie FM in a subgroup of patients or are present at concentrations too low to be detected in our assays in individuals with mild disease.

When considering the heterogeneity of FM symptoms and severity it is unsurprising that not all individuals have elevated levels of anti-SGC IgG. While pain is a uniting factor between individuals, the severity of pain and the cognitive and mood symptoms can differ greatly. Numerous studies have clustered FM patients based on pain reports, PPTs, and cognitive and psychological questionnaires (29–32). Additionally, 60% of FM patients have reduced intraepidermal nerve fiber density *(i.e.* reduced skin innervation), which is associated with more intense pain and more severe disease (4). Together these studies suggest that multiple overlapping disease mechanisms contribute to FM pathology and targeting these mechanisms may prove beneficial therapeutically. To date, questionnaire-based grouping of patients has yielded limited insight for personalized treatment. Furthermore, widespread adoption of quantitative sensory testing, intraepidermal nerve fiber density and microneurography are limited by time and skill constraints. Our study suggests that FM patients could be stratified based on autoantibody detection, which could be used for more personalized treatment approaches.

Autoantibody status is used as an indicator of disease subtype and to inform treatment strategies in other autoimmune and autoantibody-mediated diseases. For example, rheumatoid arthritis is divided into seronegative and seropositive cases where seropositive individuals have rheumatoid factor, anti-citrullinated protein antibodies or both, and seronegative individuals have neither. Seropositive rheumatoid arthritis is associated with more severe joint destruction and seropositive individuals respond better to B-cell depletion (33). Myositis disease subsets are also characterized by specific autoantibodies, which improve myositis diagnosis and help stratify treatment approaches (34–36). Our results suggest that FM patients could be divided into seropositive and seronegative subsets, creating the possibility for personalized medicine strategies. To make this a reality, future studies will need to investigate whether anti-SGC antibodies are associated with the cognitive and mood symptoms of FM and FM autoantibody targets.

FM was long thought of as a central nervous system disease, but it is increasingly clear that both the peripheral nervous system and the immune system are important contributors to FM pathology. Hyperactivity of primary nociceptors (5) and reduced intraepidermal nerve fiber density (4) in FM patients suggest that neuroplasticity occurs in the DRG, rather than only the brain and spinal cord. Both nociceptor hyperactivity and reduced intraepidermal nerve fiber density are induced by transferring FM IgG into mice (9). FM IgG also binds antigens expressed in mouse and human DRGs, further suggesting causal roles of the peripheral nervous system and immune system in FM. Peripheral immune cells, such as NK cells (37) and mast cells (38), have also been linked to FM. Our current study found anti-SGC IgG levels are elevated in individuals with more severe pain, increased pressure pain sensitivity and higher FIQ scores, but are unrelated to CPM. The lack of association with CPM suggests that anti-SGC antibodies are driving peripheral characteristics of FM. Thus, the subgroup of FM patients with anti-SGC antibodies may correspond to a group with more severe peripheral FM characteristics like nociceptor hyperactivity and reduced intraepidermal nerve fiber density.

Individuals included in both FM cohorts often had FM for many years prior to blood collection and FM diagnoses are often delayed from the disease onset (39). The timing between onset and blood collection raises the question of whether anti-SGC antibodies are an outcome of FM or a possible contributor to disease onset and persistence. FM IgG injection into mice induces FM characteristics, suggesting that autoreactive IgG contributes to FM onset (9). Here, we found that anti-SGC antibody levels are associated with FM severity but not with FM duration nor pain duration. To determine the time frame in which FM autoantibodies develop and how autoantibody levels fluctuate with time longitudinal studies are needed. Regardless, our findings indicate that FM autoantibodies are not a result of chronic disease, but instead a contributor to the disease mechanism.

In the current study we did not find a difference between FM IgG and HC IgG binding to neurons in culture. We had previously reported elevated IgG binding to sensory neurons *in vitro* (9), but these experiments were done using pooled IgG preparations. When looking at the data points from single individuals in the current study, many individuals have no or low levels of neuron binding IgG, but there are also individuals that have higher levels of IgG binding to neurons. It remains possible that a rare subset of FM patients has anti-neuron antibodies. Regardless, if autoantibodies binding SGCs are driving nociceptor hypersensitivity, then they must do so by indirectly activating SGCs or by activating neuronally expressed Fcγ-receptors (11,40).

Our assay may lack the sensitivity to detect low levels of pathogenic IgG in FM samples and it cannot differentiate between of pathogenic anti-SGC IgG and non-pathogenic, non-specific IgG binding to SGCs. For example, seven HC samples had relatively high (>50% of SGCs bound by IgG) IgG binding to SGCs. We reason that among pain-free individuals, with normal variation in their antibody repertoires, some may have IgG that binds SGCs. For example, in these individuals a recent infection could have triggered higher levels of non-specific antibodies. Similarly, serum from OA patients had IgG that bound SGCs with greater intensity compared to HC serum, but there was no difference in the percentage of cells bound. While the difference between OA and HC serum was far less pronounced than FM and HC, limited to a single parameter (intensity), and FM serum had greater levels of anti-SGC antibodies than OA serum, it is possible that some individuals with OA have low levels of anti-SGC antibodies. Although FM was an exclusion criterion in our OA subjects, up to 25% of OA patients have comorbid FM (8), so it would be interesting to examine levels of anti-SGC antibodies in subjects with severe OA with and without severe FM. Ultimately, identifying the autoantigenic targets of FM IgG will increase specificity to differentiate between FM patients with and without autoantibodies, and between other pain conditions and HC subjects that have non-pathogenic anti-SGC IgG.

Our use of murine cell cultures to assess levels of anti-SGC and anti-neuron antibodies creates a potential caveat when relating our findings to humans and FM. Mouse SGCs could express antigens that FM IgG binds, but are not expressed by human SGCs, and vice versa. However, serum IgG binding to SGCs in human DRG tissue sections is elevated and correlated with the level of anti-SGC IgG detected in cultures. Human SGCs are transcriptionally similar to mouse SGCs according to single cell RNA-sequencing studies (preprint) (41), as are human DRG neurons to mouse neurons (42), suggesting that SGCs and neurons will have similar cell surface protein repertoires. Considering that FM IgG binds both human and mouse SGCs, it is unlikely that FM IgG is binding different or at least completely unrelated targets between species.

We found that anti-SGC IgG levels are elevated in two distinct FM cohorts. The level of these antibodies correlated with a more severe clinical presentation, while being unrelated to FM duration. Moreover, the level of anti-SGC antibodies detected in culture is correlated with binding to human SGCs. These results, combined with our previous findings that FM IgG induces pain-like behaviour in mice, provide a possible answer to one mechanism of nociplastic pain in FM. Taken together, the data suggest that testing FM patients for elevated levels of anti-SGC IgG may be a relevant strategy to stratify patients for therapies that interfere with antibody function such as abatacept, anti-FcRn antibodies, IVIg or plasmapheresis.

## Author Contributions

Conception and design of the study: E Krock, E Kosek, CIS; Acquisition and analysis of data: E Krock, CEMU, JM, MH, AS, DK, JT, VV, KK, L.H., CBM, LD, E Kosek, CIS; Drafting a significant portion of the manuscript or figures: E Krock, CEMU, JM, E Kosek, CIS.

